# Self-contained and modular structured illumination microscope

**DOI:** 10.1101/2021.02.25.432746

**Authors:** Robin Van den Eynde, Wim Vandenberg, Siewert Hugelier, Arno Bouwens, Johan Hofkens, Marcel Müller, Peter Dedecker

## Abstract

We present a modular implementation of structured illumination microscopy (SIM) that is fast, largely self-contained and that can be added onto existing fluorescence microscopes. Our instrument, which we call HIT-SIM, can theoretically deliver well over 50 super-resolved images per second and is readily compatible with existing acquisition software packages. We provide a full technical package consisting of schematics, a list of components and an alignment scheme that provides detailed specifications and assembly instructions. We illustrate the performance of the instrument by imaging optically large samples containing sequence-specifically stained DNA fragments.

## 1 Introduction

Fluorescence microscopy is a proven tool to study molecules, organelles, cells and whole organisms within their native context. Super-resolution (SR) techniques elaborate on this by providing spatial resolutions that extend beyond the diffraction limit. A range of different SR methodologies are available, largely differing in their achievable spatial and temporal resolutions. Structured illumination microscopy (SIM) achieves an up to twofold increase in spatial resolution by acquiring multiple fluorescence images using different illumination patterns, followed by computational fusion of these images into a single SR image. Two-dimensional SIM requires nine fluorescence images per reconstruction, allowing it to deliver fast imaging that is theoretically compatible with any type of labeling [1]. Commercial SIM systems require between 60 ms and 1 second to acquire a single reconstructed image, while custom-built instruments exist that can deliver a full image within 20 ms [2–4]. Recent work has also shown that this can be extended to simultaneous three-dimensional acquisitions [5, 6]. The speed and label-compatibility of SIM are major advantages over slower SR techniques such as PALM/STORM/SOFI [7–10], though the latter techniques may offer considerably higher spatial resolutions. These advantages make SIM especially suited to investigations that require information on systems that are highly dynamic [11–13].

One area where SIM holds high promise is high-throughput imaging. A range of high-throughput and high-content imaging systems are readily available commercially, though high-throughput imaging at the SR level remains challenging despite multiple advances [14–19]. The fast imaging rate of SIM renders it exceptionally promising in this regard [20], though the main challenges are in the high demands placed on the instrument and on the data processing pipeline. In principle, the technique requires only standard wide-field observation and illumination of the sample using periodic patterns at different orientations and phases. However, achieving these patterns with a sufficient quality is non-trivial and requires considerable expertise in the design and alignment of optical instruments. The acquired images must also be carefully processed to deliver a reliable super-resolved image. Fortunately, well-performing open source software for SIM reconstruction is available through fairSIM [21], though identifying hardware that achieves a high performance and can be integrated in more complex measurement schemes is more challenging [3, 4, 22, 23].

In this contribution we describe a novel SIM instrument that is specifically tailored to robust and fast imaging, which we call ‘HIT-SIM’ (HIgh-Throughput Structured Illumination Microscopy). The instrument delivers fast and high-quality SIM imaging and possesses several features that render it attractive for automated scanning of large, optically thin samples. We provide schematics, software, and assembly and operating instructions in the supplementary information [24], and have devised the instrument so that key components (e.g., camera and light source) can be exchanged with modest effort. The hardware design is based on commercial components apart from a small number of custom elements that pose no particular manufacturing difficulties. Furthermore, the electronics of the system have been developed in such a way that the SIM functionality operates independently from the image acquisition software. We showcase our HIT-SIM as a fast setup to identify bacterial species based on labeled DNA, using a methodology known as ‘Fluorocode’ [25, 26].

## 2 Methods

### 2.1 Bead measurements

200 nm tetraspeck beads (Thermofisher - T7280) were used to verify the performance of the instrument. 10 μl of the beads was deposited in a 1:4 dilution (in MilliQ) on 35 mm glass-bottom microdishes (MatTek). The dish was allowed to dry overnight at 4°C in an inverted orientation. The dish was imaged using a 1.75× order selection aperture, a 488 nm 200 mW Oxxius laser with a 50 ms exposure setting, or a 561 nm 300 mW Oxxius laser with a 50 ms exposure setting on a PCO edge 4.2 camera. SIM images were reconstructed using fairSIM with the following settings: import: background subtraction 100, parameter estimation: OTF NA 1.2, lambda 525/600, a=0.3, exponential approximation, run parameter fit w/o further changes, reconstruction: Wiener 0.02, APO cut 1.9, APO bend 1, OTF attenuation off.

### 2.2 Fluorocode measurements

Bacteriophage lambda DNA was enzymatically labeled at 5’-TCGA-3’ sites using M.TaqI methyltransferase with a rhodamine B functionalized cysteine AdoMet analog as described in Ref. [26]. The slide was imaged using the 1.75× order selection aperature with a 561 nm 300 mW Oxxius laser (set to 300 mW) and 200 ms exposure setting on a PCO edge 4.2 camera. The following fairSIM settings were used for reconstruction. Import: background subtraction 100, parameter estimation: OTF NA 1.2, lambda 600, a=0.3, exponential approximation, run parameter fit w/o further changes, reconstruction: Wiener 0.05, APO cut 2, APO bend 0.9, OTF attenuation off.

## 3 Results and Discussion

### 3.1 Optical system

The design of the HIT-SIM microscope is based on the two-beam fastSIM microscope reported in Ref. [2]. A schematic representation of the system is shown in figure 1. The system consists of a conventional wide-field microscope that can generate periodic illumination patterns using a spatial light modulator (SLM). The SLM and other alignment-sensitive SIM-specific illumination components are combined into a single mechanically-rigid ‘SIM module’ (figure 1 and figure 2).

**Figure 1:**
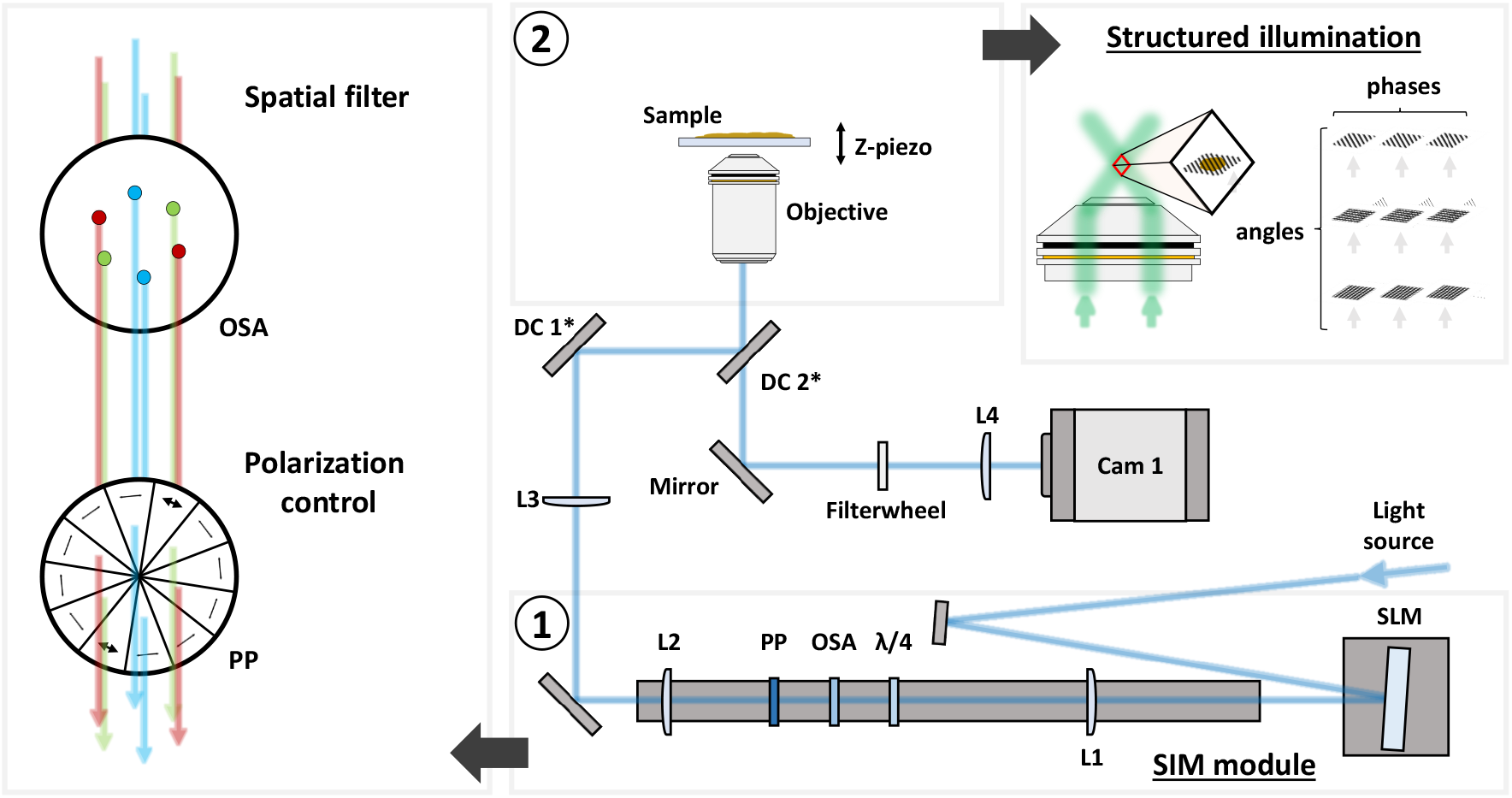
Schematic representation of the HIT-SIM. With ➀ the SIM module, ➁ a representation of the structured illumination. L:lens, OSA:Order Selection Aperture, λ/4: Quarter waveplate, PP:Pizza Polarizer, DC:Dichroic, SLM:Spatial Light Modulator.

**Figure 2:**
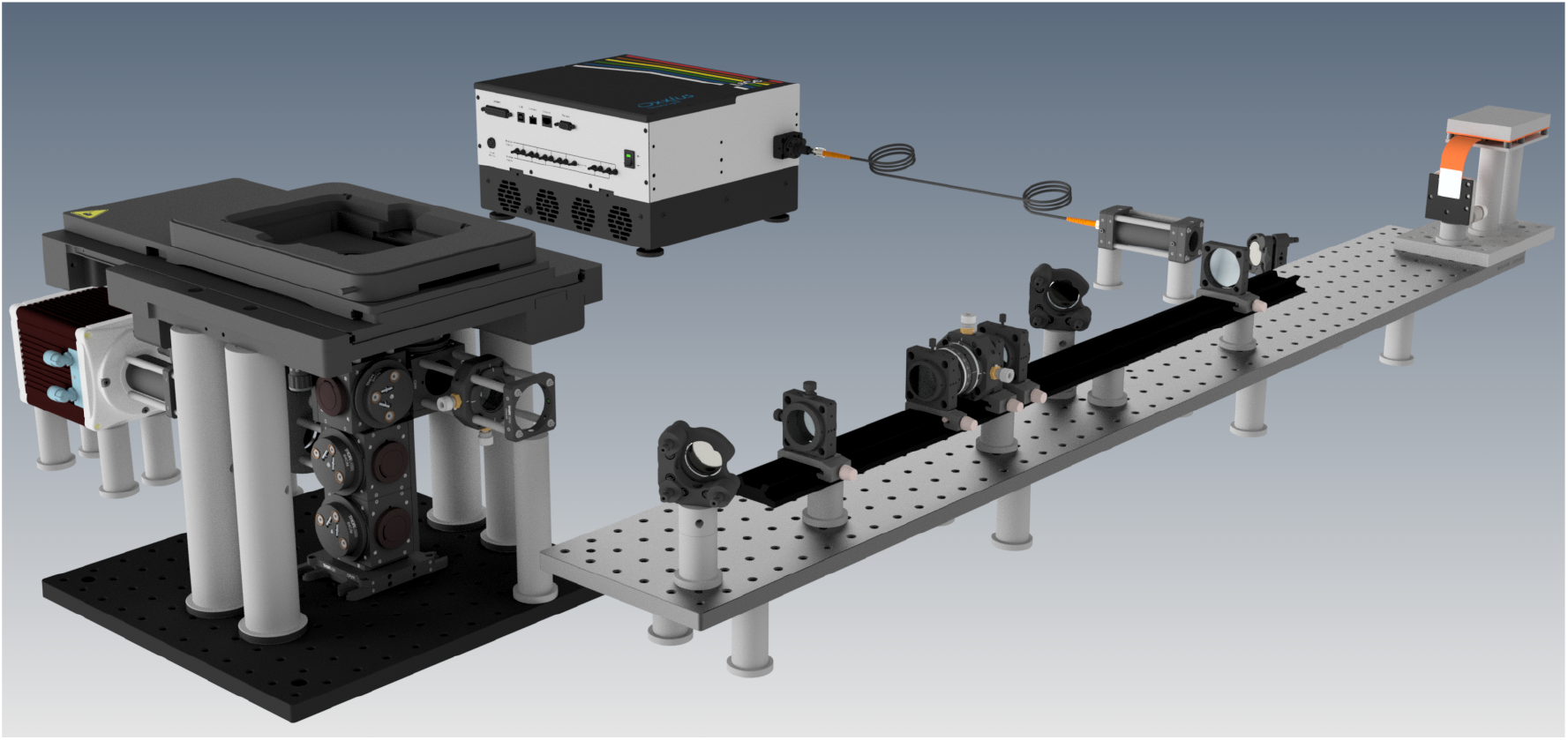
CAD rendering of the HIT-SIM system.

Light from the laser source is directed onto the SLM, which acts as a configurable diffraction grating that can switch between patterns in less than 500 μs. These SLM patterns are chosen such that they yield two first order beams that produce a sinusoidal illumination profile at the sample. Stray diffraction orders, which arise due to the binary nature of the SLM, are excluded from the excitation path by focusing the light via lens L1 onto a metal disk with precisely drilled holes, the order selection aperture (OSA). The choice of the focal length of L1 depends on the pixel size of the SLM, the desired range of resolution enhancements and wavelengths, and the size constraints of the downstream optics, as is discussed in the supplementary information [24] and accompanying documents. We also require precise control of the excitation polarization [27, 28], which is accomplished using a quarter waveplate (QWP, *λ*/4) and segmented ‘pizza’ polarizer (PP). The ratio of the focal lengths of lenses L2 and L3 are chosen to match the pattern with the desired resolution for the objective in use.

The SLM pattern generation was carried out with a multi-color search algorithm [3], which itself is based on previous work for SLM-based SIM [2, 28]. In practice, one may want to define several sets of SLM patterns and OSA configurations that provide different spacings of the illumination pattern, allowing a trade-off between the achievable resolution increase, z-sectioning capability, and sensitivity to abberations and noise. Changing to a different resolution enhancement then requires an exchange of the OSA and selecting a different SLM pattern set in software. The SLM patterns can be also chosen such that a single ‘color independent’ OSA can be used for all design wavelengths, providing a uniform resolution enhancement factor. We opted for a single mask that accepted both 488 and 561 nm illumination.

The excitation light is reflected into the objective using two dichroic mirrors, DC1 and DC2. The use of a single dichroic mirror would lead to a degradation of the SIM pattern since the *s* and *p* polarizations are subject to different phase shifts upon reflection from a typical commercial dichroic. This can be corrected by reflecting the light from two identical dichroic mirrors positioned at orthogonal angles [23] (DC1 and DC2; supplementary information [24]). The collected emission light is transmitted through DC2 and focused on the camera using lens L4, which was chosen to yield a ≈100 nm pixelsize. The field of view in our implementation is approximately 90×90 μm when combined with a 60× water immersion objective.

The instrument can acquire a full SIM image stack in well under 50 ms. Assuming an exposure time of 1 ms, SLM changeover time of ≈1.9 ms and a camera readout time of ≈2.5 ms, the HIT-SIM can grant an imaging speed of 25 reconstructed SIM images per second, though in practical experiments the acquisition speed is usually limited by the available laser power, noise levels, and the concentration and brightness of the fluorophores in the sample.

### 3.2 Instrument control

Operating the instrument requires multiple levels of control and analysis to (i) synchronize the various components during the acquisition of a SIM image, (ii) perform imaging experiments involving automated acquisitions at different points on the sample, and (iii) reconstruct the SIM images from the raw fluorescence images.

At the lowest level, the SIM acquisition requires sub-millisecond synchronization of the excitation light sources, SLM, and camera. Such precise timing requirements are difficult or impossible to achieve using software running on general-purpose operating systems, and are also not required in the vast majority of wide-field based imaging experiments. As a result, there are usually no provisions for such synchronization in image acquisition software, nor are there off-the-shelf solutions to embed this functionality into existing imaging pipelines. The usual solution is to implement custom logic into a separate microcontroller combined with the development of custom acquisition software to control this device as well as the other required hardware (cameras, light sources). This custom solution is then tied to the particulars of the instrument in question, and often does not contain more advanced functionality such as the automated imaging of larger samples.

While the use of dedicated hardware such as a microcontroller is unavoidable, we reasoned that many of the resulting limitations could be mitigated if the measurement software does not need to be aware that the instrument is performing a SIM measurement. In fact, this distinction comes naturally with SIM, since detection of the fluorescence requires only classical wide-field imaging. For single-color acquisitions, there is no need for the acquisition software to even be aware of the SLM or microcontroller, as long as it is capable of replacing the acquisition of a single fluorescence image with the acquisition of nine consecutive fluorescence images. The microcontroller, in turn, is responsible only for synchronizing the SLM and light source with the camera exposure, but need not be aware of any other hardware, nor does it need to communicate with the acquisition computer. The only requirement is that it can detect when the camera begins exposing, and that it knows that each SIM acquisition will consist of nine fluorescence images in total, where the displayed SLM pattern must be advanced between every camera acquisition.

In practice, a suitable synchronization signal is readily provided by the exposure output that is present on all scientific cameras, and that typically transitions to logic high upon the start of a camera exposure, regardless of how this exposure was initiated. This signal remains high for the full duration of the camera exposure, during which the microcontroller enables the light source and synchronizes its output with the SLM. The microcontroller detects the end of the exposure by monitoring for a transition back to logic low while the image is being read out, activating the next SLM pattern in time for the start of the next acquisition.

In our implementation, the microcontroller monitors this signal and assumes that the start of a camera exposure means that a SIM acquisition is being initiated. It will then automatically illuminate the sample with the first SIM pattern, managing the laser illumination and SLM functionality until the exposure signal transitions back to logic low. If the camera initiates another exposure within 500 ms, the microcontroller assumes that we wish to measure the next SIM pattern, which it will display for the duration of this subsequent exposure. This procedure loops over the full set of nine SLM patterns, resetting back to the beginning of the sequence after nine consecutive exposures have been detected, or at any time when there has been no new exposure within 500 ms of the previous one. The measurement software does not need to be aware that a SIM measurement is running, and is free to perform fluorescence experiments using the full range of functionality required by the end user. Our strategy is also compatible with multi-color imaging, where the microcontroller must be aware that the different excitation wavelengths will always be acquired in the same sequence (e.g., nine fluorescence images with green excitation, followed by nine fluorescence images with red excitation). A more detailed description of this strategy is given in the supplementary information [24].

While any camera control software can be used in practice, we decided to use the Micro-Manager software [29] for the implementation of the system detailed here. In addition to its openness and broad range of device support, Micro-Manager also provides direct integration of the SIM reconstruction software available through the fairSIM project [21]. Thanks to our design, no changes need to be made to the software to enable the acquisition of SIM images, and the full Micro-Manager functionality is readily available.

### 3.3 Imaging performance of the HIT-SIM instrument

We verified the performance of our system by dual color imaging of fluorescent beads. Figure 4 shows 200 nm beads imaged with 488 nm and 561 nm excitation light using an OSA and SLM pattern set designed for a 1.75× improvement in resolution (for the 488 and 561 nm lasers respectively, an expected resolution of 116.2 and 133.6 nm). Here, as anticipated, individual beads can be clearly resolved from the SIM reconstruction while being unresolvable in conventional wide-field imaging (approximated here by summing the nine raw fluorescence images). The Fourier transforms of these SIM images display the typical pattern intrinsic to SIM (supplementary information [24]), supporting the enhanced spatial information. As expected, the achieved resolution of the 561 nm illuminated beads is a factor 561/488 worse than that of the 488 nm illuminated beads due to the longer wavelength of the light.

**Figure 3:**
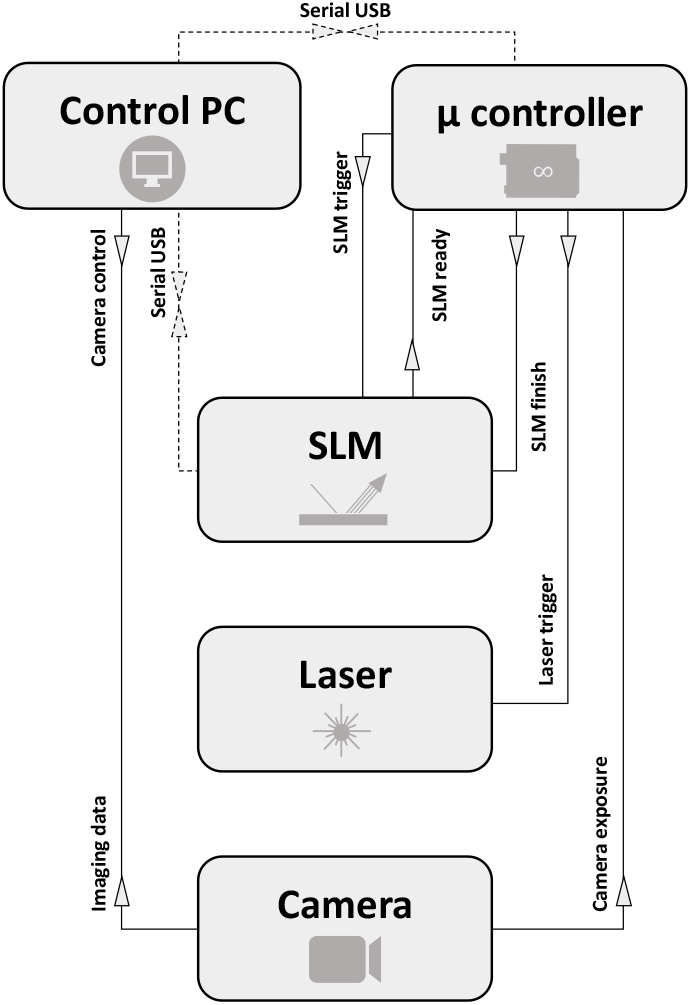
Representation of the HIT-SIM electronics triggering scheme that allows for a simple exchange of modules. The dotted connections are only required for setting up/configuring the system.

**Figure 4:**
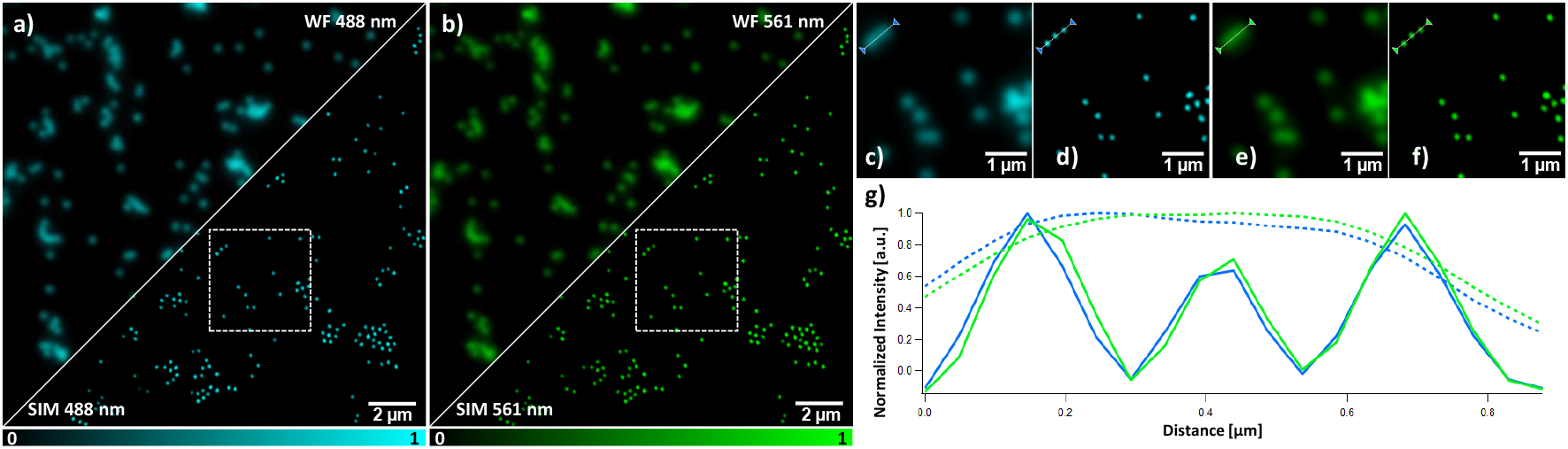
200 nm tetraspeck beads imaged with HIT-SIM in two colors. (a) wide-field (above the cut) versus SIM (below the cut) with 488 nm excitation. (b) wide-field (above the cut) versus SIM (below the cut) with 561 nm excitation. (c) Wide-field and (d) SIM of the inset in (a), 488 nm excitation. (e) Wide-field and (f) SIM of the inset in (b), 561 nm excitation. (g) Lineplot of the region marked in (c), (d), (e) and (f), with wide-field in dotted lines and SIM in full lines (488 and 561 nm excitation in blue and green respectively). This dataset was obtained using the 1.75× enhancement settings.

We found that our instrument can reach imaging speeds up to 30.43 mm^2^/hour. The majority of this time is in fact spent waiting for the sample stage position to stabilize after a move command (750 ms per field-of-view; supplementary information [24]). The overall imaging duration can be subdivided into the actual image acquisition (13.74%), active XYZ stage movement (18.90%), and waiting for stage stabilization (67.36%). These results suggest that a more performant choice of stage could greatly speed up these acquisitions.

As a final validation we applied the HIT-SIM instrument to the automated scanning of large samples, exemplified here by a sample loaded with large DNA molecules that are sequence-specifically labeled using the ‘Fluorocode’ methodology [25, 26] (figure 5). The resulting fluorescence distribution is determined by the sequence of each DNA fragment and can be used to identify the organism that produced the fragments. The achievable accuracy of this is limited by, among others, the spatial resolution with which the fluorophores can be positioned along the DNA fragment and the number of fragments that can be read out.

**Figure 5:**
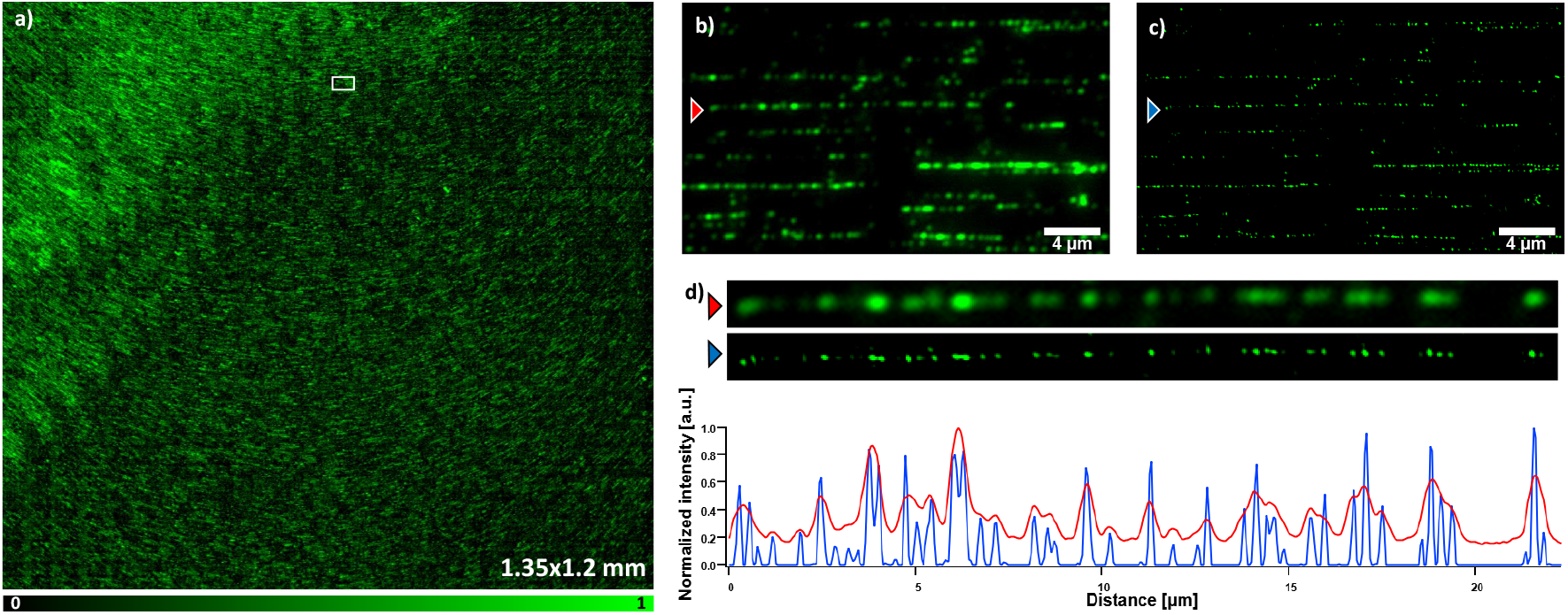
Fluorocode imaging with HIT-SIM. (a) Stitched wide-field image (displayed 1.35×1.2 mm), with an inset magnified in (b) and (c). (b) Wide-field and (c) reconstructed SIM with respectively red and blue arrows indicating a labeled DNA sequence. A zoom of the DNA sequence from (b) and (c) accompanied by a line plot (respectively red and blue) are given in (d). This imaging sequence was obtained using the 1.75× resolution enhancement OSA and 561 nm laser.

We reasoned that the combination of high required spatial resolution and large sample area were well suited to the fast and automated imaging of our instrument, taking advantage also from the fact that our software design allows direct usage of the high-content imaging support in Micro-Manager. We upgraded our measurement using automatic focusing by interpolation of the focus position between manually-focused reference positions on the sample, though our SIM module is also compatible with solutions for active focus control based on the reflection of infrared light. We imaged a sample consisting of bacteriophage lambda DNA labeled at 5’-TCGA-3’ sites with a rhodamine B functionalized cysteine AdoMet analog, acquiring an area of 1.59×1.5 mm with a 1.75× improvement in spatial resolution compared to conventional imaging. This enabled us to create a fully stitched image (of which 1.35×1.2 mm is shown in 5) using in-house software, showing the strong improvement in information made possible by using our instrument.

## 4 Conclusion

The HIT-SIM instrument is a robust and fully documented implementation for fast SIM imaging, designed to function as a self-contained module both in optical layout and in instrument control. This design choice provides compatibility with a wide range of components, requiring only an exposure output from the camera and trigger input to the light source, and can work with virtually any image acquisition software. The optical performance of our instrument was validated by both measurement of samples consisting of fluorescent beads and samples consisting of sequence-specific labeled DNA. The use of a fast switching SLM provides support for the imaging of fast dynamic processes such as mitochondrial fusion/fission or viral transfer, but also for the imaging of extended sample regions. We expect that our system can considerably accelerate the applications of SIM in high-throughput or high-content settings.

## Acknowledgements

The authors would like to thank Viola Mönkemöller for assistance with the measurements and data analysis, and Laurens D’Huys for assistance with the Fluorocode sample. R.V. and S.H. thank the Research Foundation Flanders for a doctoral and postdoctoral fellowship, respectively. P.D. thanks the Research Foundation Flanders for grants G0B8817N and G090819N, the KU Leuven for grant C14/17/111, and the European Research Council for grant 714688 NanoCellActivity. M.M. thanks the European Commission for a Marie Skłodowska-Curie individual fellowship. J.H. acknowledges financial support from the Horizon 2020 Framework Programme of the European Union ADgut (grant number 686271); from the European Union Research Council through the ERC-2017-PoC Metamapper (grant number768826); from the FWO (Fonds voor Wetenschappelijk Onderzoek, grant number G0C1821N); from the Flemish government through long term structural funding Methusalem (CASAS2, Meth/15/04) and from the Max Planck Institute as MPI fellow.

